# Microcosm cultures of a complex synthetic community reveal ecology and genetics of gut microbial organization

**DOI:** 10.1101/2022.09.13.507837

**Authors:** Xiaofan Jin, Feiqiao B. Yu, Jia Yan, Allison Weakley, Katherine S. Pollard

## Abstract

The behavior of microbial communities depends on both taxonomic composition and physical structure. Metagenomic sequencing of fecal samples has revealed the composition of human gut microbiomes, but we remain less familiar with the spatial organization of microbes between regions such as lumen and mucosa, as well as the microbial genes that regulate this organization. To discover the determinants of spatial organization in the gut, we simulate mucosal colonization over time using an *in vitro* culture approach incorporating mucin hydrogel microcosms with a complex yet defined community of 123 human strains for which we generated high-quality genome assemblies. Tracking strain abundance longitudinally using shotgun metagenomic measurements, we observe distinct and strain-specific spatial organization in our cultures with strains enriched on mucin microcosms versus in supernatant, reminiscent of mucosa versus lumen enrichment *in vivo*. Our high taxonomic resolution data enables a comprehensive search for microbial genes that underlie this spatial organization. We identify gene families positively associated with microcosm-enrichment, including several known for biofilm and adhesion functions such as efflux pumps, gene expression regulation, and membrane proteases, as well as a novel link between a coenzyme F420 hydrogenase gene family and lipo/exopolysaccharide biosynthesis. Our strain-resolved abundance measurements also demonstrate that incorporation of microcosms yields a more diverse community than liquid-only culture by allowing co-existence of closely related strains. Altogether these findings demonstrate that microcosm culture with synthetic communities can effectively simulate lumen versus mucosal regions in the gut, providing measurements of microbial organization with high taxonomic resolution to enable identification of specific bacterial genes and functions associated with spatial structure.

## Main

Human gut microbiomes consist of diverse microbial taxa [1, 2], with typical complexity ranging on the order of over a hundred species in a single individual [3]. Spatial organization of gut microbes is linked to community function and host health [4–10] – in particular, different taxa are enriched between mucosa and lumen [11–16], and mucosal colonizing bacteria may be particularly able to regulate host-microbiome interactions and immunomodulation [17–21]. However, we still lack a high-taxonomic-resolution view of ecological differences between lumen and mucosa, and accordingly possess a limited understanding of genetic factors underlying this spatial structure. As within-species dynamics exist within gut microbiomes [22–25], we hypothesize that distinct spatial organization may (i) occur at the level of individual strains, and (ii) be associated with specific gene families and pathways that regulate mucosa versus lumen colonization.

To test our hypotheses, we develop an integrated experimental-computational workflow that compares lumen- and mucosal-like niches within a complex gut community. By using metagenomic sequencing, we are able to profile microbes with high taxonomic resolution, enabling strain- and gene-level analysis. We use a synthetic 123 strain community modeled closely after the recently published hCom2 community [26, 27], cultured *in vitro* with added mucin microcosms to provide a mucosal-like substrate for bacterial attachment distinct from the surrounding liquid supernatant [28, 29]. To identify genetic correlates of microcosm colonization, we develop a computational workflow that uses a comprehensive search across KEGG Orthology (KO) gene families [30] to identify associations between gut spatial organization and underlying microbial genotypes, using phylogenetic regression to account for evolutionary relationships between taxa [31–33].

Our approach provides key advantages over existing alternatives: first, by using an in vitro approach that allows mucin microcosm and supernatant subpopulations to be independently sampled [29] – analogous to mucosa and lumen *in vivo* – we obtain information on spatial structure missing from stool sampling and traditional liquid culture. Independent sampling of lumen and mucosal subpopulations is also possible using *in vivo* human gut biopsy, but the invasiveness of this approach limits sample sizes and longitudinal measurements [34]. In contrast, our *in vitro* platform enables us to sample mucosal- and lumen-like community subpopulations across multiple passage timepoints, with statistical replicates. Second, using our defined 123-strain community – which we generate high quality genomes for each member therein – allows us to emulate the bacterial complexity found in human guts, yet still accurately quantify abundance using metagenomic sequencing even between closely related strains. Strain-level measurements are critical for enabling gene-level analysis, as they allow genetic comparisons between closely related taxa. By comparison, earlier work with microcosms used 16S sequencing of undefined communities to produce measurements with more limited taxonomic resolution and did not examine genes associated with microcosm colonization [29].

We demonstrate that this approach yields detailed strain-level measurements of differential spatial organization, revealing taxa which are reproducibly enriched or depleted on mucin microcosms relative to supernatant. Then, we identify numerous genes and biosynthetic gene clusters that distinguish microcosm versus supernatant genomes consistently across phylogenetic lineages, including genes related to cell adhesion and biofilm formation whose presence differs between closely related strains with distinct microcosm enrichment profiles.

## Results

### Closed genomes enable strain-level metagenomic profiling of complex defined microbial communities

Starting from isolate cultures of 123 bacterial strains that are prevalent in the human gut microbiome (Fig. 1A, Table S1), we first generate high-quality, contiguous genomes for all strains other than five with closed genomes already. For the other 118 strains, we perform hybrid assembly of long Nanopore (median 3.9 × 10^4^ reads/strain) and short Illumina reads (median 1.7 × 10^6^ reads/strain) (Fig. S1, Methods), successfully generating closed assemblies with no more than 10 contigs. By contrast, the closest available NCBI genome (Fig. 1A) is more fragmented (78/123 comprise more than 10 contigs) and less closely related to the strain in our defined community; 20/123 have *>* 0.1% ANI difference to our strain, and 33/123 contain 100 or more differential KEGG Orthology (KO) gene families (see Methods, Fig. S2). Thus, our reference database of closed genomes that are exact strain matches is critical for accurate strain and gene-level characterization of metagenomic data. Next, isolate strains are combined into a single community using anaerobic automated liquid handling (See Methods, Fig. S3), and inoculated into cultures containing 0.5% mucin 1% agar microcosms and MEGA media with 6 3-day passages (Fig. 1B, Methods). As a control, we also culture in parallel the same inoculum with MEGA media only, i.e., liquid-only culture. We use metagenomic sequencing of microcosms and supernatant sampled independently (1.2 × 10^7^ read pairs per sample) at each passage to quantify strain relative abundances (see Methods, Fig. S4). To analyze read libraries with high taxonomic resolution, we use NinjaMap [27] with our custom genome database to generate strain-level abundances (Fig. 1C, Table S2) – we successfully validate NinjaMap results against lower taxonomic resolution species-level abundances generated using Kraken2 [35] with the UHGG database [2] (median *R*^2^ = 0.978070 across samples, see Fig. S5).

**Figure 1:**
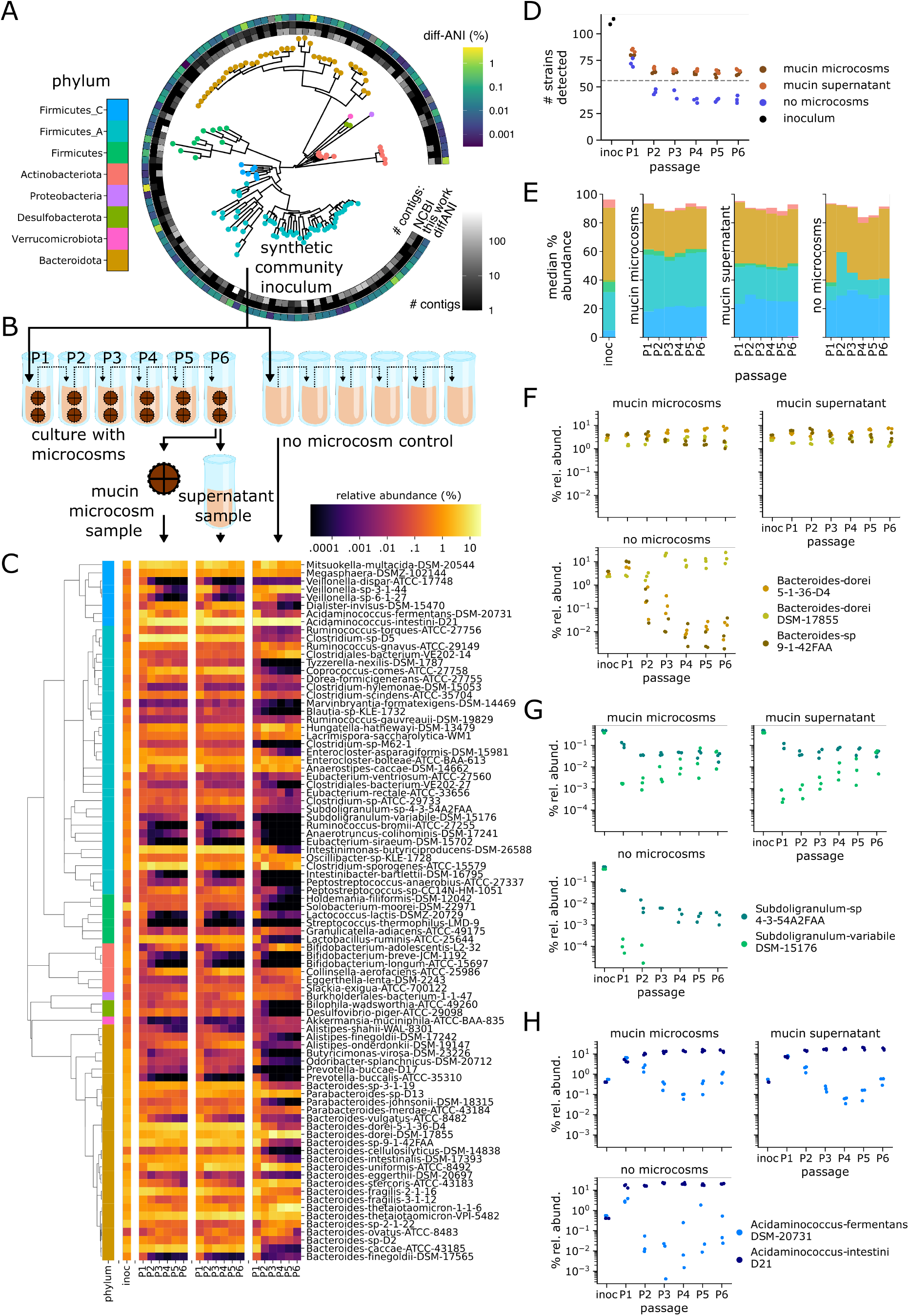
Microcosm cultures yield stable, diverse communities with coexistence of closely related strains. **(A)** We generate closed, high quality genomes for each strain in a 123-member microbial community, representative of taxa in the human gut. De novo generated genomes are more contiguous than closest previously available NCBI genomes, and represent exact matches to our strains. **(B)** We use this 123-member community to inoculate cultures incorporating mucin microcosms as well as non-microcosm controls. We passage (P) each culture 6 times (3 days between passages), independently sampling bacterial DNA from microcosm, supernatant, and no-microcosm control at each timepoint for downstream metagenomic sequencing. **(C)** We use NinjaMap to obtain community relative abundances from metagenomic sequencing data, here we plot median abundance of each strain at each passage timepoint, across experimental conditions. **(D)** Number of detected strains after culture stabilization (~ P3 and later) is higher in microcosm versus no microcosm cultures, indicating enhanced community richness. Grey dashed line indicates median number of strains detected (56) using same threshold with the 119-member hCom2 community in mice [26] **(E)** Addition of microcosms leads to broad taxonomic shifts in community composition relative to no-microcosm control, visualized here at phylum level. **(F)** Strain-resolved abundance patterns of 3 B. dorei strains (*ANI >* 99%) in our community demonstrates stable co-existence enabled by addition of microcosms, compared with dominance of a single *B. dorei* strain without microcosms. **(G)** Strain-resolved abundance patterns of 2 *Subdoligranulum* strains (*ANI* ~ 80%) in our community demonstrates stable co-existence enabled by addition of microcosms. *Subdoligranulum variabile* DSM-15176 in particular also exhibits increasing abundance over passage timepoints. **(H)** Strain-resolved abundance patterns of 2 *Acidaminococcus* strains (*ANI* ~ 80%) demonstrates more stable co-existence when cultured in the presence of microcosms.

### Mucin microcosms increase community richness and promote strain-coexistence within *in vitro* cultures

Next, we characterize differences that result from spatial structure introduced by the incorporation of mucin microcosms. In cultures without microcosms, community richness drops from a median of 113.5/123 detected strains in the inoculum (detection cutoff 0.01% relative abundance, 1% horizontal coverage – some strains with non-viable glycerol stocks / isolates did not grow to sufficient ODs, see Fig. S7), stabilizing down to a median of 38.5/123 detected strains by the 4 late passages (passages P2-6, i.e., days 9-18). By contrast, microcosm cultures seeded with the same inoculum stabilize to a median of 62.5/123 and 65.5/123 detectable strains on microcosms and supernatant respectively (Fig. 1D). This significantly elevated richness ((*p <* 1^*−*5^), see Fig. S6) parallels results from hCom2 inoculated in mice (median 56/119 detected strains across 19 mice) [26], suggesting cultures with microcosms provide a closer analog to *in vivo* conditions than do liquid-only cultures. Increased richness is particularly noticeable in Firmicutes, Firmicutes_A, and Bacteroidota (see Fig. S6), while total abundance is higher for Firmicutes and Firmicutes_A, but lower in Bacteroidota (Fig. 1E).

Beyond phylum level effects, abundance shifts also occur at strain level. Addition of microcosms increases abundance for a diverse set of strains including *Bacteroides caccae* ATCC-43185, *Lactobacillus ruminis* ATCC-25644, *Coprococcus comes* ATCC-27758 (which displays extremely sticky / slime phenotype in pure culture), two strains in family Marinifilaceae (*Butyricimonas virosa* DSM-23226, *Odoribacter splanchnicus DSM-20712*), and both sulfur reducing bacteria (*Desulfovibrio piger* ATCC-29098 and *Bilophila wadsworthia ATCC-49260* from phylum Desulfobacterota). Some taxa are largely unaffected by microcosms, such as three Bifidobacterium strains, while few taxa are negatively affected by microcosms, with three closely related Veillonella strains being notable exceptions (see Fig. S8). These strain-level abundance shifts do not always align with corresponding phylum-level shifts, emphasizing the value of our highly-resolved taxonomic measurements.

One of the most striking abundance shifts revealed by strain-level analysis is the co-existence of closely related strains with the addition of microcosms. In liquid-only culture, *Bacteroides dorei* DSM-17855 outcompetes two closely related (ANI *>* 99%) strains, *Bacteroides dorei* 5-1-36-D4 and *Bacteroides sp*. 9-1-42FAA (Fig. 1F). By contrast, these three strains coexist stably in culture when microcosms are present. Other examples can be found between two closely related (ANI ~ 80%) Firmicutes_A strains: *Subdoligranulum sp*. 4-3-54A2FAA and *Subdoligranulum variabile* DSM-15176 (Fig. 1G), and between two closely related (ANI ~ 80%) Firmicutes_C strains: *Acidaminococcus fermentans* DSM-20731 and *Acidaminococcus intestini* D21 (Fig. 1H). These observations of co-existence (see Fig. S8 for additional examples) concur with increased richness detected in microcosm cultures.

To better understand why community richness increases with mucin microcosms, we additionally grow our inoculum in cultures with 1% agar microcosms (plain-agar, i.e., no mucin). We observe that plain-agar microcosm culture also exhibits overall enhanced richness compared with liquid-only culture (Fig. S6). However, we do note some specific strain-level differences between mucin-agar and plain-agar microcosm cultures (See Fig. S7,S8). This suggests that increased richness result largely – but not entirely – from having a physical surface to colonize rather than the nutrients provided by mucin. These results are reminiscent of similar effects in bacterial biofilms, where increased diversity has been attributed to expanded spatial niches and reduced competition [36–38].

### Strains exhibit distinct enrichment profiles between microcosm and supernatant communities

We next characterize spatial organization *within* microcosm cultures by comparing subpopulations sampled from microcosm and supernatant, testing our hypothesis that strain-level spatial differences occur within gut communities. For the 86/123 prevalent strains that are detected in at least 10% of passaged samples (see Methods), we quantify a microcosm enrichment score – defined as the log fold change in abundance between paired microcosm and supernatant samples (i.e., derived from the same culture tube) – for each strain and each passage (Fig. 2A, Table S3). We also calculate a single aggregate, normalized log-microcosm-enrichment score for each strain based on late passage measurements (see Methods). These scores reflect the preference of each strain to grow on microcosms versus in the supernatant, with positive scores indicating microcosm preference.

**Figure 2:**
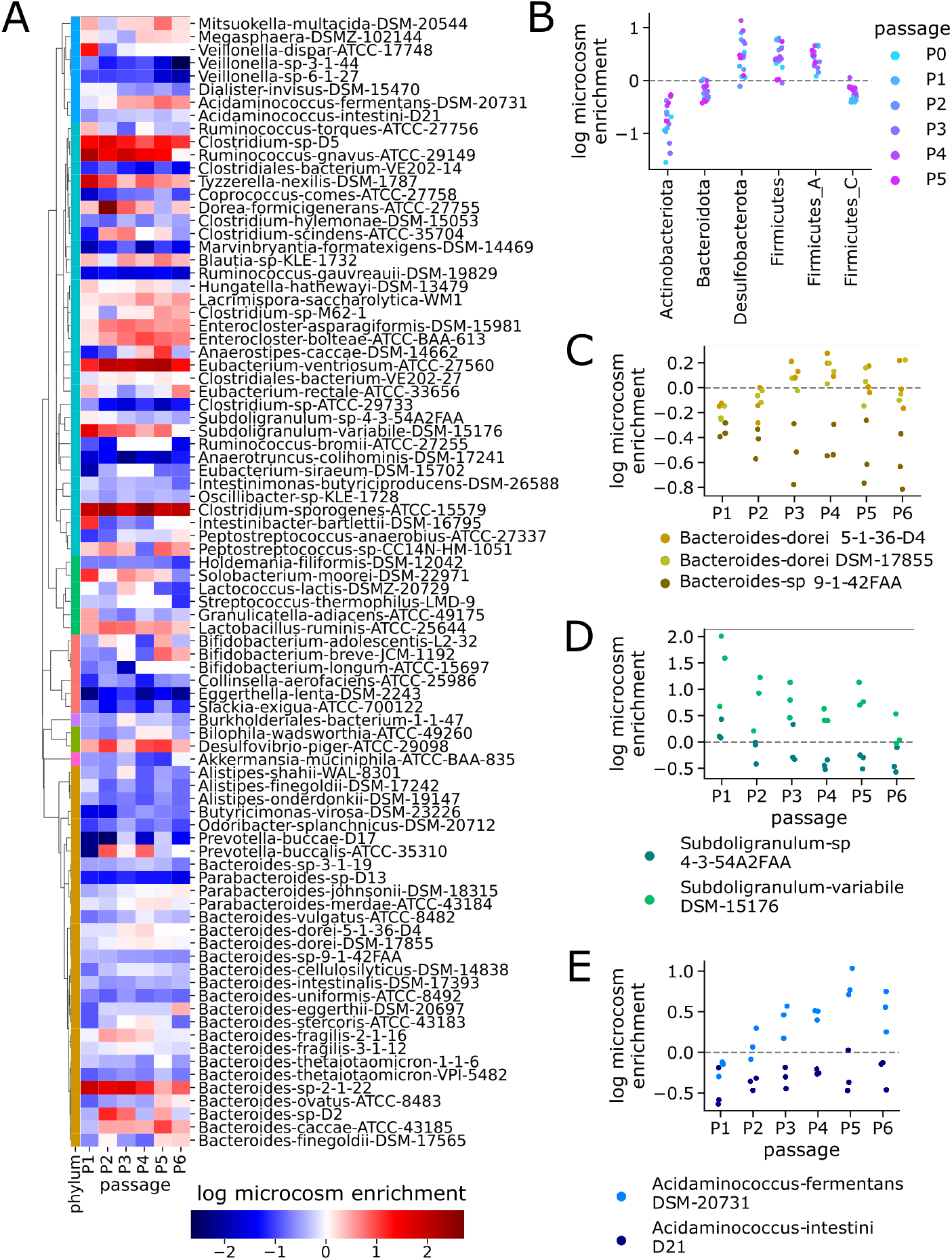
Strain level differences exist between mucin microcosm and supernatant communities. **(A)** Strains exhibit different microcosm enrichment phenotypes, both within and between clades – positive (red) scores indicate higher relative abundance on microcosms versus supernatant. **(B)** Aggregated at phylum level, taxa exhibit evidence of distinct spatial structure: Desulfobacterota, Firmicutes and Firmicutes_A are enriched on microcosms, while Actinobacteriota, Bacteroidota and Firmicutes_C are enriched in supernatant. **(C)** One of the three *Bacteroides dorei* strains (sp. 9-1-42FAA) exhibits consistent microcosm depletion relative to the other two strains (5-1-36-D4 and DSM-17855). **(D)** *Subdoligranulum variabile* DSM-15176 exhibits consistent microcosm enrichment relative to the closely related strain *Subdoligranulum sp*. 4-3-54A2FAA. **(E)** *Acidaminococcus fermentans* DSM-20731 exhibits consistent microcosm enrichment relative to the closely related strain *Acidaminococcus intestini* D21. Additionally, microcosm enrichment of *Acidaminococcus fermentans* DSM-20731 increases with time towards later passages.

Aggregating at phylum level, we observe enrichment toward mucin microcosms in Desulfobacterota, Firmicutes (primarily Bacillus-like), and Firmicutes_A (primarily Clostridia-like), and enrichment toward supernatant in Actinobacteriota, Bacteroidota, and Firmicutes_C (primarily Negativicutes-like), with no obvious time-dependent signal (Fig. 2B). These results are largely consistent between mucin-agar and plain-agar microcosms, with the exception of Desulfobacterota which is not enriched on plain-agar microcosms. Certain individual strains also exhibit similar trends, such as Eubacterium ventriosum ATCC-27560 which exhibits microcosm preference with mucin-agar microcosms but not plain-agar (Fig. S9).

At strain level, we find a diverse range of enrichment profiles over time (Fig 2A), including several strains with opposite enrichment relative to their phylum. For instance, *Bacteroides sp*. 2-1-22 prefers microcosms, while *Clostridiales bacterium* VE-202-14 from phylum Firmicutes_A prefers supernatant. Moreover, closely related strains can exhibit different enrichment phenotypes: *Bacteroides dorei* 5-1-36-D4 and DSM-17855 exhibit similar abundance in supernatant and microcosm (log enrichment scores *≈* 0), but *Bacteroides sp*. 9-1-42FAA displays consistent enrichment toward supernatant (log enrichment scores *<* 0, see Fig. 2C). *Subdoligranulum variabile* DSM-15176 and *Acidaminococcus fermentans* DSM-20731 prefer mucin microcosms more than their respective counterparts, *Subdoligranulum sp*. 4-3-54A2FAA (Fig. 2D) and *Acidaminococcus intestini* D21 (Fig. 2E). These findings support our hypothesis that distinct strain-level spatial organization occurs within gut communities.

Finally, as external validation we compare our in vitro microcosm enrichment results against an *in vivo* dataset [15] with paired mucosal and lumen samples (see Methods, Table S6). We find our *in vitro* microcosm-enrichment strain scores exhibit similarity to *in vivo* mucosal-enrichment species scores within inter-subject variability (Fig. S10, Table S7). We also observe general agreement at phylum level: Bacteroidota is enriched toward both supernatant *in vitro* and lumen *in vivo*, while Firmicutes_A and Firmicutes are enriched toward microcosm / mucosa. However, discrepancies also exist, as Actinobacteriota is enriched toward supernatant *in vitro* and mucosa *in vivo* (Fig. S10). These results suggest that our experimental platform provides a close – though not exact – approximation of *in vivo* structure.

### Phylogenetic regression predicts genes associated with mucosal colonization

We next test for statistical associations between microcosm enrichment and underlying microbial genotypes, evaluating our hypothesis that key microbial genes may regulate spatial organization in the gut. Using kofamscan [30] to comprehensively search all genomes against all defined KO families, we generate a genotype matrix consisting of 9857 KOs detected in the 86 prevalent strains. Each entry in this 86 × 9857 matrix corresponds to maximum kofamscan/hmmer bitscore hit for a particular KO in a particular genome (Fig. 3A) – higher scores reflect gene presence. We then test for each of the 9857 KOs whether its genotype pattern across the 86 strains is significantly associated with the corresponding pattern of microcosm enrichment scores (phenotype). We perform significance tests using phylogenetic regression with phylolm [32] to account for evolutionary relationships between strains (see Methods).

**Figure 3:**
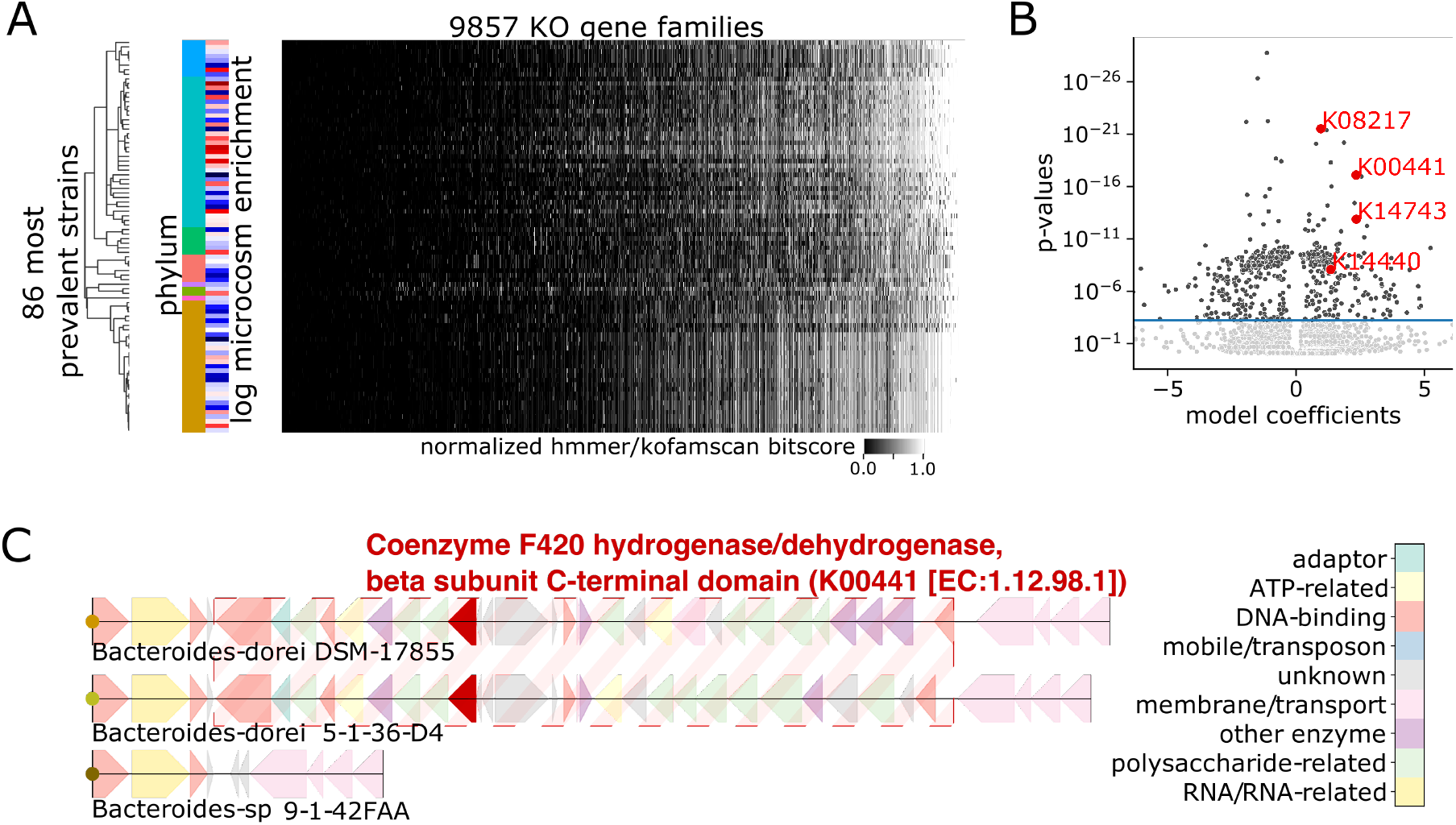
Phylogenetic regression identifies genes associated with mucin microcosm enrichment. **(A)** Phylogenetic regression identifies significant associations between log microcosm enrichment score (red/blue indicates positive/negative microcosm enrichment respectively) and gene presence absence patterns (lighter/darker shades of gray indicate gene presence/absence respectively) across the most prevalent 86 strains detected in passaged samples. We use this model to test a total of 9857 KEGG KO gene families determined using kofamscan [30], accounting for phylogenetic relatedness between strains assuming Brownian motion along evolutionary branches. **(B)** Volcano plot of phylogenetic regression test, where each dot represents one KEGG KO – horizontal line at FDR=0.01. Horizontal axis is clipped at 0.1 and 99.9 percentiles, highlighted gene families colored in red. **(C)** *Bacteroides dorei* 5-1-36-D4 and DSM-17855 both harbor a coenzyme F420 dehydrogenase gene (KEGG KO K00441, *hmmerbitscore* = 190.7, 188.7) colocalized amongst LPS/EPS related gene clusters – these features are collectively missing from the corresponding region in the *Bacteroides sp*. 9-1-42FAA genome.

Our approach identifies 244 KO families significantly associated with increased enrichment on mucin microcosms relative to supernatant applying FDR correction at p<0.01 threshold (Fig. 3B, see also Methods, Table S8). Out of these KOs, we highlight several illustrative examples whose genotype patterns align with differential microcosm enrichment in the *B. dorei, Subdoligranulum* and *Acidaminococcus* strains featured in Fig. 2C-E. From the three *B. dorei* strains, we find two KO gene families in particular – K00441 (coenzyme F420 hydrogenase subunit beta [EC:1.12.98.1], Fig. 3C, 4A) and K08217 (MFS transporter, DHA3 family, macrolide efflux protein, Fig. 4B) – which have strong homology hits in *Bacteroides dorei* 5-1-36-D4 and DSM-17855, but not in *Bacteroides sp*. 9-1-42FAA. Mapping the K00441 coenzyme F420 hydrogenase hits to their genomic loci in 5-1-36-D4 and DSM-17855, we find the gene resides in the midst of lipo/exopolysaccharide (LPS/EPS) biosynthesis gene clusters (Fig. 3C). Performing gene neighborhood analysis across all 123 strain genomes to search for KOs enriched within 10 kilobases (kb) of K00441 annotated genes, we find 107 hits (see Methods, Table S5), which are dominated by KOs with LPS/EPS biosynthesis functions including numerous glycosyltransferase, epimerase, sugar-reductase, polysaccharide membrane transporter genes, suggesting a previously uncharacterized link between coenzyme F420 hydrogenase and microbial LPS/EPS production.

**Figure 4:**
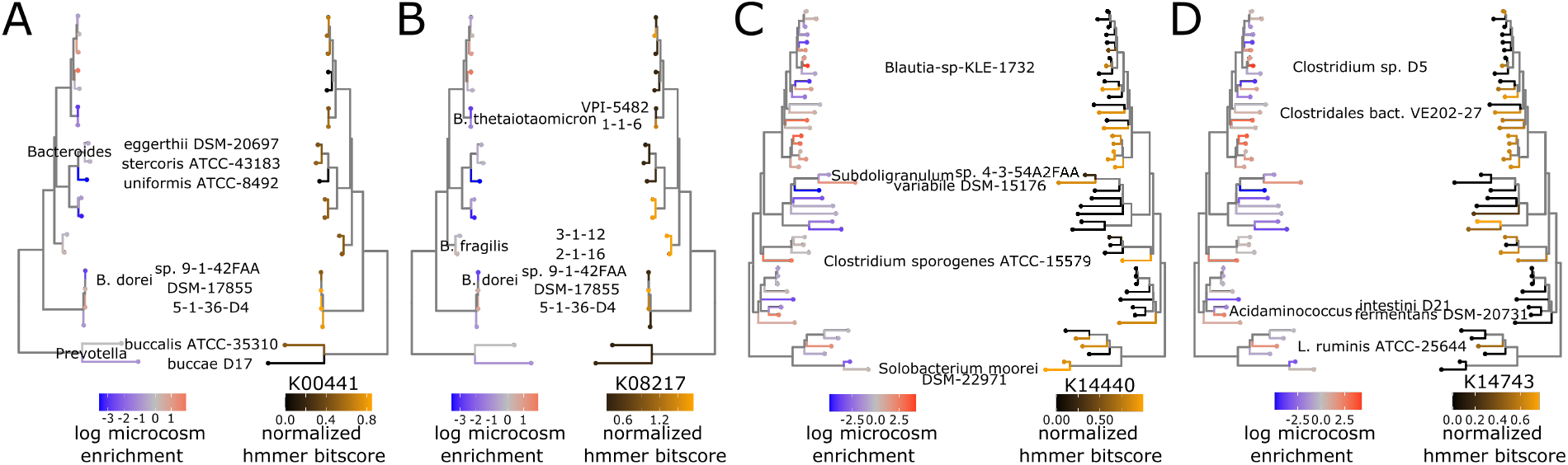
Phylolm-identified gene families have presence patterns that align with differential microcosm enrichment. **(A)** Comparison of microcosm enrichment pattern (left) with gene presence pattern (right) of K00441 coenzyme F420 hydrogenase subunit beta [EC:1.12.98.1], across family Bacteroidaceae strains. **(B)** Comparison of microcosm enrichment pattern (left) with gene presence pattern (right) of K08217 MFS transporter, DHA3 family, macrolide efflux protein, across family Bacteroidaceae strains. **(C)** Comparison of microcosm enrichment pattern (left) with gene presence pattern (right) of K14743 membrane-anchored mycosin MYCP [EC:3.4.21.-], across phylum Firmicutes_A, Firmicutes_C, and Firmicutes strains. **(D)** Comparison of microcosm enrichment pattern (left) with gene presence pattern (right) of K14440 SWI/SNF-related matrix-associated actin-dependent regulator of chromatin subfamily A-like protein 1 [EC:5.6.2.-], across phylum Firmicutes_A, Firmicutes_C, and Firmicutes strains.

Beside K00441 and K08217 in *B. dorei*, we also note a strong hit to a DEAD box helicase gene family – K14440, SWI/SNF-related matrix-associated actin-dependent regulator of chromatin subfamily A-like protein 1 [EC:3.6.4.12] – in *Subdoligranulum variabile* DSM-15176 (Fig. 4C) and a membrane protease gene family – K14743, membrane-anchored mycosin MYCP [EC:3.4.21.-] – in *Acidaminococcus fermentans* DSM-20731 (Fig. 4D) which are absent in their less microcosm-enriched relatives. Intriguingly, LPS/EPS biosynthesis [39–43], membrane transporters/efflux pumps [44–50], membrane proteases [51–56], and DEAD box helicase gene regulators [41,57–60] all have known links to biofilm formation and adhesion. Aggregating all 244 microcosm-associated KOs by KEGG BRITE gene categories, we identify several BRITE categories enriched for significant KOs, representing antibiotic resistance genes, glycosyltranferases (E.C. 2.4), phosphotransferases (E.C. 2.7), transcriptional regulators, and proteases (Table S11), further supporting the importance of these gene functions in mucosal colonization.

Testing for clade-specific effects using within-phylum phylogenetic regression, we find K14440 and K14743 to be among the most significant hits in Firmicutes/Firmicutes_A/Firmicutes_C, while K00441, K14743 and K08217 are among the most significant hits for Bacteroidota (Table S9). As an external validation, we repeat our workflow using the Suez *et. al in vivo* dataset [15] to identify a list of KOs associated with mucosal enrichment (Table S10), and find statistically significant overlap between genes associated with microcosm enrichment *in vitro* and genes associated with mucosal enrichment *in vivo*, (*log − odds − ratio* = 4.0, *p* = 9.7 × 10^*−*21^, Fig. S11). Thus, we confirm that measurements from our *in vitro* synthetic community cultures are sufficiently detailed to inform a computational gene-level analysis of gut spatial organization, revealing that genes related to biofilm formation and adhesion likely play key roles in modulating gut microbial structure.

### Strain enrichment on microcosms is associated with presence of lipo/exopolysaccharide biosynthesis gene clusters

To explore mechanisms of community structure beyond individual genes, we next investigate microcosm enrichment of biosynthetic gene clusters (BGCs). We use deepBGC [61] to search for BGCs across our strain genomes, annotate BGCs based on their KEGG KO presence, and apply hierarchical clustering to categorize 1103 detected BGCs into 256 groups with similar KO co-occurrence patterns (Fig. 5A, Table S12). We then map presence/absence of each of these 256 BGC-groups against the 86 prevalent strains in our experiment (Fig. S12), and apply phylogenetic regression to test for associations between microcosm enrichment and BGC-groups.

**Figure 5:**
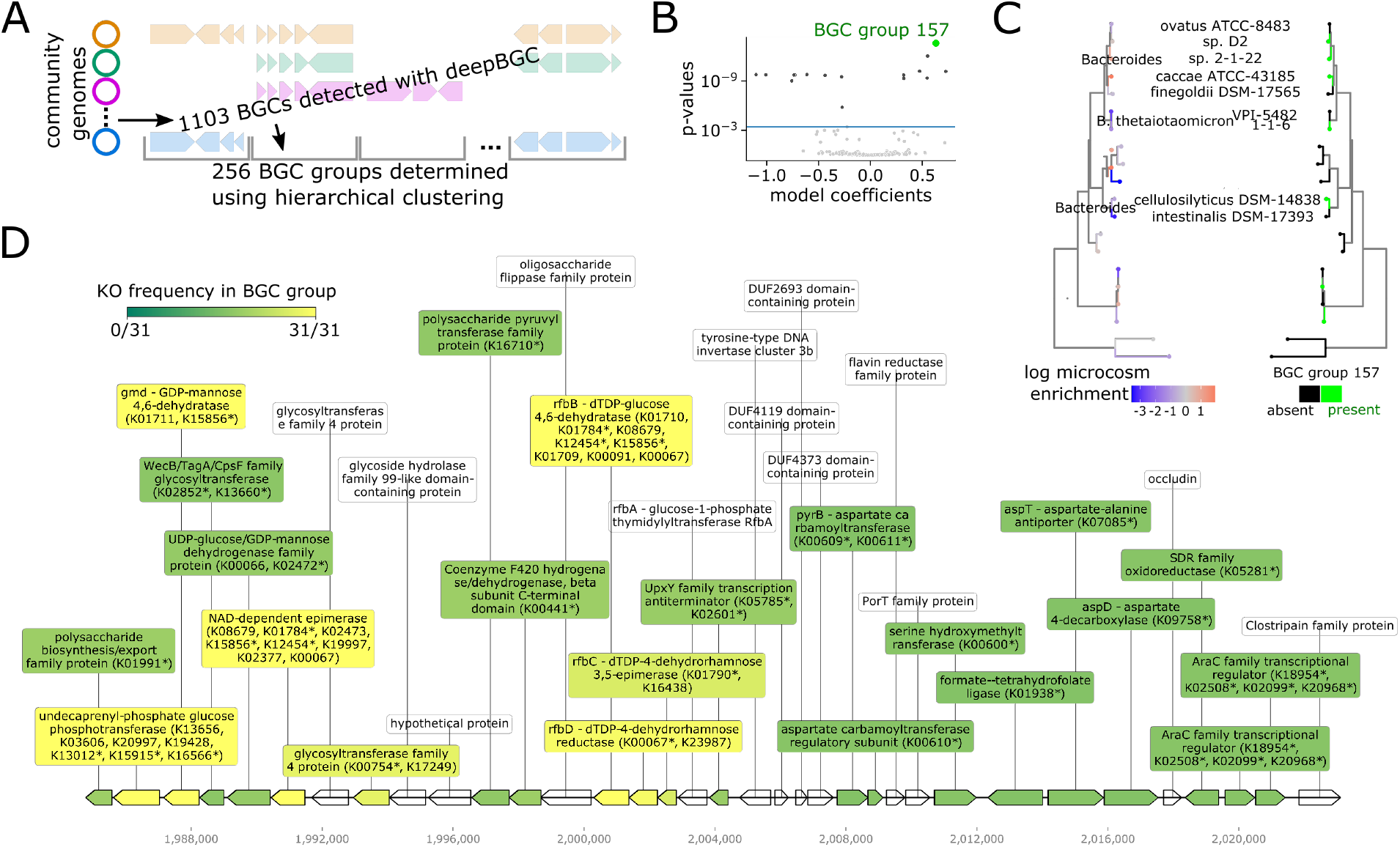
Strain enrichment on mucin microcosms is associated with exopolysaccharide gene clusters. **(A)** Schematic of approach used to generate and group BGCs across strains using deep-BGC and hierarchical clustering. **(B)** Volcano plot of phylogenetic regression test, each dot represents one BGC-group, horizontal line at FDR=0.01 cutoff. Top hit BGC-group 157 highlighted in green. **(C)** Comparison of microcosm enrichment pattern (left) with presence pattern (right) of BGC-group 157 across strains in family Bacteroidaceae. **(D)** Example of a representative gene cluster in BGC-group 157 from the *Bacteroides* strain with highest microcosm enrichment score (*Bacteroides-sp*.*2-1-22_cluster_1_1984776-2023109*.*1*). Gene label colors reflect frequency of KO family among all BGCs in group – in cases where a gene maps to multiple KOs, * marks mapped KO with highest frequency in BGC-group.

Our approach yields a total of 7/256 significant BGC-groups positively associated with microcosm enrichment (Fig. 5B), the three largest of which consist of 18 or more BGC representatives (BGC-group 157 – see Fig. 5C, BGC-group 120, and BGC-group 69). Filtering for the most common KEGG KOs in each of these BGC-groups, we discover that BGC-group 157 and BGC-group 120 consist of likely EPS related gene clusters, typified by glycosyltransferase, epimerase and other EPS related KOs (Fig 5D, Table S13). BGC-group 69 consists largely of gene clusters populated by membrane transporter genes. KOs in other microcosm enriched BGC-groups include more polysaccharide related genes (BGC-groups 198, 186, 161) and AraC transcriptional regulator genes (BGC-group 34). These findings at the BGC-level further reinforce our KO-level results, showing that membrane-related functions such as LPS/EPS and transporters, as well as key gene regulators, likely regulate spatial organization in our *in vitro* model of the human gut.

## Discussion

Applying mucin microcosm culture with our defined community of human gut strains, we present here the first strain-resolved measurements of spatial structure within the context of a complex gut microbial community. By *a priori* generating a database of high quality closed reference genomes, our approach enables high taxonomic resolution abundance measurements using metagenomic sequencing, while effectively recapitulating spatial structure in the gut microbiome. These measurements show with high taxonomic resolution how a complex gut microbial community is spatially organized upon introduction of microcosms, demonstrating that microcosms enhance community richness to a level similar to *in vivo* observations, including instances of co-existence between closely related strains. We find clear enrichment signals *within* microcosm cultures where certain strains prefer to grow on the microcosms versus in the supernatant, or vice versa. Microcosm enrichment phenotypes can differ significantly even between closely related strains, supporting our hypothesis that spatial organization in the gut occurs at strain-level and trends would be missed at coarser taxonomic resolution.

Another benefit of using strain-resolved metagenomics is that we can identify gene families that specifically occur in strains with microcosm enrichment (or depletion) phenotypes. We do so using phylogenetic regression, a rigorous statistical approach that adjusts for evolutionary relationships between strains. This analysis identifies several gene families related to microbial adhesion and biofilm formation, including efflux pumps (e.g., K08217) that are known to mediate collective biofilm phenotypes such as quorum sensing and antibiotic resistance [44–50], and membrane proteases (e.g., K14743) which can enhance motility / colonization on surfaces [51–56]. We also find genes involved in biosynthesis of LPS/EPS which are known to mediate bacterial adhesion [39–43], such as glycosyltransferase and epimerase genes, as well as a particular gene family K00441 (coenzyme F420 hydrogenase subunit beta [EC:1.12.98.1]) for which we report significant genomic colocalization with other known LPS/EPS genes, suggesting a previously uncharacterized functional link. We also find several groups of biosynthetic gene clusters containing membrane transporters and LPS/EPS genes associated with microcosm enrichment. Beyond membrane-associated functions, our analysis also highlights regulatory genes such as SWI/SNF DEAD box helicases (K14440). Intriguingly, such genes have not only been shown to be involved in biofilm formation [41, 57–60], but also specifically drive expression of efflux pumps and LPS/EPS genes [57]. We speculate that in mucosa-associated taxa, key regulator genes act as master switches for a host of bacterial functions that alter outer membrane composition to enhance biophysical interactions with the mucosal surface and thus increase mucosal colonization fitness, leading to global spatial organization of these taxa towards the mucosa (Fig. 6).

**Figure 6:**
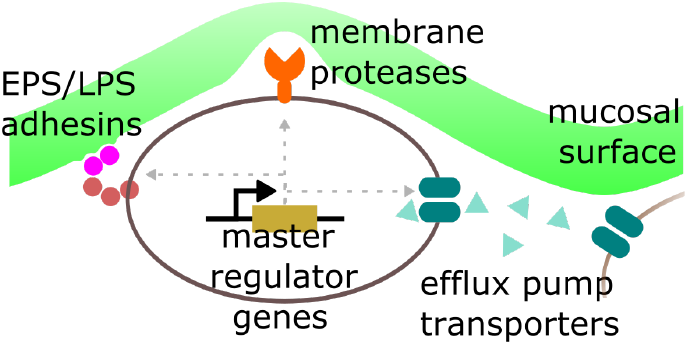
Schema for how mucosal associated genes may regulate spatial structure. Depiction of proposed framework where in mucosa-associated taxa, regulatory genes serve as master switches for microbial functions that increase mucosal colonization fitness such as LPS/EPS, membrane transporters / efflux pumps, and proteases.

We conclude by noting several limitations to our work and point to areas for further exploration. First, we only show statistical associations – not causal mechanisms – between genotypes and microcosm enrichment, meaning hits should be cautiously interpreted as potential genetic factors deserving of followup investigation. Synthetic biology in genetically tractable gut strains can be used to test our predictions by altering the expression of identified gene families using gene knockout, knockdown or knockin experiments [62–64]. Second, while our *in vitro* results generally parallel those from earlier *in vivo* work [15, 26], we do find limited discrepancies (e.g., microcosm depletion of Actinobacteriota), meaning our current platform provides a close but still imperfect replica of the *in vivo* gut environment. More realistic culture conditions can be explored, potentially through modification of media conditions (e.g., addition of bile acids, different carbon sources). Third, our current approach based on metagenomic sequencing provides accurate quantification of strain and gene abundance, but it does not assay gene expression or spatial localization on microcosms. Future work using gut microbial metatranscriptomic analysis [65, 66] and multiplexed FISH imaging [67–69] can greatly complement current capabilities and mitigate these shortcomings. Fourth, it remains unclear how strain-strain interactions affect structure. Follow-on studies with our platform that incorporate strain dropout can address these questions. Finally, in addition to strains from healthy Western guts, future work should incorporate taxa found in dysbiotic and non-Western guts to explore how spatial structure varies between healthy and diseased states, and across global geographic regions. Ultimately we believe the platform presented here has the potential to transform the standard for *in vitro* investigation of gut microbiota, in a manner that recognizes the important interplay between spatial structure and strain-level ecology.

## Methods

### Hybrid assembly of microbial isolates

Strains are cultured in isolation until stationary phase, followed by DNA extraction using phenol chloroform. DNA is sequenced using both Oxford Nanopore long-read and Illumina short-read sequencing, followed by hybrid assembly using custom bioinformatic workflow (Fig. S2) built using Unicycler [70], RScaf [71] and TGS-GapCloser [72] – workflow is available as docker images, see Software availability below.

### Community phylogeny

Phylogenetic tree structure of the community is generated using GTDB-tk [73], using our genome assemblies as input.

### Genome annotation and gene classification

Genomes are annotated using NCBI PGAP [74]. Predicted protein sequences are then mapped using kofamscan [30] to the to KEGG Orthology database.

### Mucin microcosm preparation

Mucin microcosms are prepared similarly to previously described protocols [28, 75], using boiled 0.5% porcine mucin (Sigma M2378) and 1% agar (BD 214030) solution solidified onto K1 biofilm carriers (Evolution Aqua MEDIAK1). Mucin free agar-only microcosms are prepared using 1% agar solution.

### *In vitro* culture of synthetic community with mucin microcosms

To construct the full *in vitro* synthetic community, we first culture each strain in isolation in 1.8 mL of its preferred media in a 96 well deep well plate (Table S1). Because of the large range of growth rates and stationary phase cell densities, strains are inoculated in a staggered fashion with slow growers inoculated 3 days prior and fast growers inoculated 1 day before community assembly. Fastidious growers are cultured in 10 mL and concentrated to increase final cell density. Individual isolate cultures are sequenced to verify purity. On the day of community assembly, cell density for each strain is estimated using OD measured on a plate reader (BioTek Epoch). Using this measurement, each strain is normalized to a maximum OD of 0.3 using liquid handling robotics. Cultures are pelleted and washed with PBS, and then combined to form a mixture of 123 strains (epMotion 5073). Strains are combined in an anaerobic environment equipped with automated liquid handling in order to reduce potential cross contamination and other human errors when concurrently handling many strains (Fig. S3).

Following assembly of our bacterial community, the mixture is used to inoculate cultures in 15 mL tubes comprising MEGA media and 5 microcosms each. Cultures are left to grow at 37*°C* in anaerobic conditions for 3 days without agitation, at which point they are passaged. Passaging consists of transferring a single microcosm from the old culture tube to a new culture tube. This process is repeated 5 times for a total of 6 passages - for each subsequent passage, the previously transferred-in microcosm is discarded prior to transferring of a microcosm to the next culture. For liquid-only cultures, inoculating loops (Fisherbrand 01-189-165) are used for passaging. Supernatant pellets and microcosm samples are saved and frozen at each passage point.

For each condition, we culture the community in biological triplicate cultures (i.e. 3 separate culture tubes). Each culture tube is sampled with technical triplicates – for microcosm samples, we pick 3 microcosms out of each culture tube to store at *−*80*°C* prior to DNA extraction, while for supernatant and liquid-only cultures, we take 3 separate 1mL aliquots from each tube, pellet, then store at *−*80*°C* prior to DNA extraction. This yields a total of 9 read libraries for each passage and experimental condition. The initial inoculum communities are sampled in duplicate, each sample sequenced 3 times each.

### DNA extraction, Library prep and sequencing

DNA is extracted from pellets and microcosms using ZymoBIOMICS 96 DNA Kit and bead beating with 0.1mm glass beads (Benchmark Scientific D1031-01). Extracted DNA from each sample is quantified in 384 well plates on a fluorescent plate reader (BioTek Neo2) using the Quant-iT PicoGreen assay (ThermoFisher). To generate input DNA for our high-thoughput and low-volume Nextera XT library preparation process, DNA samples are normalized to at maximum of 0.2 ng/uL in a 384 well plate using a low volume cherry picking liquid handler (SPT). Library preparation is done in 384 well plates using a low-volume 16 channel liquid handler (SPT) and follows the chemistry of the Nextera XT process but in a total volume of 4 uL in order to reduce library preparation cost. Libraries are quantified again using the Quant-iT PicoGreen assay and normalized. After pooling and cleaning using Ampure XP beads (Beckman), libraries are sequenced on a Novaseq 6000 (Illumina) to a mean depth of 1.2 × 10^7^ read pairs per sample. In addition to DNA derived from microbial communities, we also sequenced all input strains used to construct the community to ensure strain purity and identity.

### Read mapping and abundance estimation

Read mapping is performed with NinjaMap as previously described [27], using our de novo generated genomes as reference database. Briefly, reads are aligned to genome sequences, with only perfect unique matches considered in the first round. Ambiguous reads are held in escrow for the first round, and subsequently assigned in a statistically weighted manner determined by initial abundance estimates from the first round of alignment. This generates relative abundance and horizontal genome coverage estimates for each strain in each sample’s read library. We consider a strain present in a sample if it exceeds a 1% horizontal coverage and 0.01% relative abundance cutoff. Out of all 270 passage samples (6 passages × 5 experimental conditions – mucin-agar microcosms, mucin-agar supernatant, plain-agar microcosms, plain-agar supernatant, no-microcosms – × 9 replicates), we use a prevalence cutoff of 10% presence (i.e. present in 27 or more samples) to focus on the most prevalent strains. For down-stream abundance-related analysis, we collapse technical (i.e., within tube) triplicates to their median abundance measurements, while considering biological (different culture tubes) triplicate measurements separately. Table S2 lists relative abundance and horizontal coverage across strains, passages, replicates and experimental conditions.

### Mucosal enrichment calculations

For each strain, and passage, microcosm enrichment scores are calculated as log ratio of microcosm to supernatant abundance, for 3 biological replicates, replacing zeros with half-minimum non-zero value prior to taking log. For each strain, a single aggregate microcosm is generated by taking the mean of over mean over standard deviation of 12 log ratio scores in the late passages (P3-6, 4 passages × 3 biological replicates). Table S3 lists enrichment scores per strain.

### Gene neighborhood enrichment test

Based on results from kofamscan for each gene in each genome, a gene is annotated with a KO-label if it exhibits overlap greater than 0.5× coverage with the KO’s pHMM model, as well as a bitscore greater than 0.5× the KO’s bitscore threshold. We count the frequency of all annotated KOs within 10 kb of K00441-labeled genes across the full community genome database. To generate p-value estimate of this measured frequency, we compare it against a null distribution generated by 1000 random gene order permutations. In each of these 1000 permutations, we randomly reassign gene labels within each of the 123 genomes prior to conducting frequency counts. p<0.01 indicates 990 or more times out of 1000, the actual co-occurence of a particular KO within 10 kb of K00441 is greater than random.

### Phylogenetic regression

For each KO family, and each strain, we determine the maximum hmmer bitscore hit to the KO’s pHMM out of all the strain’s proteins. Aggregating across KOs and strains, this yields a strain-by-KO genotype matrix, where each entry is the highest bitscore value – higher bitscores indicate gene presence. We then test for association between this genotype and microcosm enrichment score (phenotype). While such genotype-phenotype tests are in many ways similar to those conducted in genome association studies (GWAS), the application of ordinary least squares (OLS) regression, often used in GWAS, is not appropriate here due to phylogenetic relationships between strains. These relationships mean that assumptions of independence between measurements inherent to OLS are violated. We confirm the presence of a non-star phylogeny between strains by generating a phylogenetic tree based on strain genomes, using bac120 multiple sequence alignment with GTDB-tk [73] (Fig. 1A). Therefore, to account for this phylogenetic relatedness, for each KO we apply phylogenetic regression to test for significant association between mucosal enrichment scores and maximum hmmer bitscore (standard scaled) across strains. We implement this test using the R package phylolm [32], assuming a Brownian motion model along evolutionary branches, using the bac120 phylogenetic tree as input. This generates effect size estimates and p-values for every KO. We filter KOs for significance applying a p<0.01 cutoff with Benjamini-Hochberg FDR correction. In addition to running phylogenetic regression across all 86 prevalent strains, we also run these models across subsets of these strains grouped by phylum to search for clade-specific hits. For this analysis, we group Firmicutes, Firmicutes_A, Firmicutes_C phyla into a single clade.

### Comparison with *in vivo* dataset

We analyze from the Suez 2018 [15] dataset all read libraries from untreated (i.e., naive) individuals, for whom lumen and mucosal reads were available from cecum, descending colon, and terminal ileum, i.e., a total of 6 read libraries per individual. We first use Kneaddata (part of the Biobakery suite [76]) to perform host (i.e. human) filtering of read sequences, resulting in 13 individuals for which all 6 libraries exceed a read depth of 10^4^ reads (Table S6). For these 13 individuals, we obtain abundance estimates at all 6 sites by mapping reads to UHGG database using Kraken2 [2, 35]. We then calculate normalized mucosal enrichment scores for each species defined as log ratio of mucosal to lumen abundance. Score are normalized by taking mean-over-standard-deviation across all individuals and sites (13 individuals × 3 sites – cecum, descending colon and terminal ileum – for 39 total measurements). We determine gene presence absence for these species, across KOs, by using kofamscan to search the UHGG pangenome database [2], and then apply phylogenetic regression as described above to test for associations between gene presence absence and mucosal enrichment score across UHGG species. The regression uses enrichment scores from 676 species detected with greater than 0.01% relative abundance in at least 10% of *in vivo* read libraries, which contain a total of 12,822 detected KEGG KO gene families (Table S7,S10).

### Extraction and grouping of biosynthetic gene clusters using DeepBGC and hierarchical clustering

DeepBGC [61] is used to extract BGCs from our de novo genomes. For each identified BGC, we generate a list of present KOs based on if contained genes map to KO’s pHMM with overlap greater than 0.5× coverage, as well as a bitscore greater than 0.5× the KO’s bitscore threshold. We filter out BGCs with fewer than 3 present KOs, and then use hierarchical clustering to cluster all remaining BGCs based on their binary KO presence/absence profile into 256 BGC-groups, applying a Jaccard distance metric. We then map presence of each BGC-group within community strains, and use this presence/absence matrix to test for associations with microcosm-enrichment applying phylogenetic regression as described above.

## Supporting information

Supplemental Tables archive

## Data availability

All sequencing data of this study is deposited in the Sequence Read Archive (SRA), accession codes pending. Genomes are deposited in Genbank (NCBI), also pending.

## Code availability

Code used for analysis and visualization available at: https://github.com/xiaofanjin/gut-community-microcosms

Software used for nanopore basecalling and hybrid assembly available at: https://github.com/czbiohub/microbiome-data-analysis/ https://github.com/FischbachLab/nf-hybridassembly

## Acknowledgements

We thank A. Lind, P. Bradley, A. Bustion, and I.H. Riedel-Kruse for helpful discussions regarding data analysis and feedback on the manuscript, A. Cheng and M. Fischbach for sharing bacterial strains, and S. Jain and X. Meng for help building the genome assembly framework. This work is supported by funding from Chan Zuckerberg Biohub, Gladstone Institutes, NSF grant #1563159.

## Author’s contributions

XJ, BY and KP contributed to the design and implementation of the research, to the analysis of the results and to the writing of the manuscript. JY and AW contributed to the implementation of the research.

