## Supplemental Information for "Microcosm cultures of a complex synthetic community reveal ecology and genetics of gut microbial organization"

### Hybrid Nanopore Illumina genome assembly approach

As described in the Methods, we use a hybrid assembly approach combining Nanopore and Illumina reads from isolate strain DNA to generate closed genomes for each strain in the community. Each strain is cultured in its preferred medium in an anaerobic environment until it reaches stationary phase to recover cell pellets. In order to preserve the length of genomic DNA during extraction, cell pellets are subjected to more gentle bead beating (10 Hz for 5 min) and enzymatic lysis (Lucigen). Because columns tend to shear genomic DNA during the extraction process, high molecular weight DNA is isolated from cell lysate using phenol chloroform approaches and pelleted using ethanol precipitation. We resuspend DNA in elution buffer (Qiagen) and quantify both length and concentration of the extracted DNA (Agilent, ThermoFisher). We try to achieve a length of $>$ 10 kbp at $>$ 200 ng per strain.

To generate both long and short read data, we use MinION (Oxford Nanopore) and NovaSeq (Illumina). Oxford Nanopore libraries are generated using the PCR-free ligation kit (LSK109). Because long read sequencing can be cost prohibitive at scale, we multiplex 4-8 strains on the same MinION flow cell using ligation barcoding expansion kits (NBD104, NBD114), yielding 200k reads with N50 of 6-9 kbp. With bacterial genome sizes of 5 Mbp, 200k Nanopore reads cover the genome more than 100X. To alleviate the increased error rate of Nanopore long reads, we supplement Nanopore long reads for each strain with 2-3 million Illumina read pairs, which achieves a short read coverage of 100X and can be obtained at $10 inclusive of library preparation. Together, our pipeline to acquire 100X fold coverage of both long Nanopore reads and short Illumina reads for all 123 strains require a cost of $200 per strain, inclusive of reagents and consumables of all experimental steps.

Hybrid assembly of strain genomes is carried out using a custom bioinformatic approach built around Unicycler (Fig. S1) [1]. Unicycler can handle as input data a maximum of 100X coverage for long reads and short reads separately. Therefore, long reads are first filtered and sub-sampled for quality and length, whereas short reads are filtered by quality and then coverage normalized. Unicycler takes as input filtered reads and, if possible, creates a closed and polished assembly. When the assembly is not closed, scaffolding using LRScaf [2] and gap closing using TGS-GapCloser [3] are performed in an attempt to generate a complete reference genome. The strain hybrid assembly pipeline using Nanopore and Illumina reads is made available as a docker image `fischbachlab/nf-hybridassembly`.

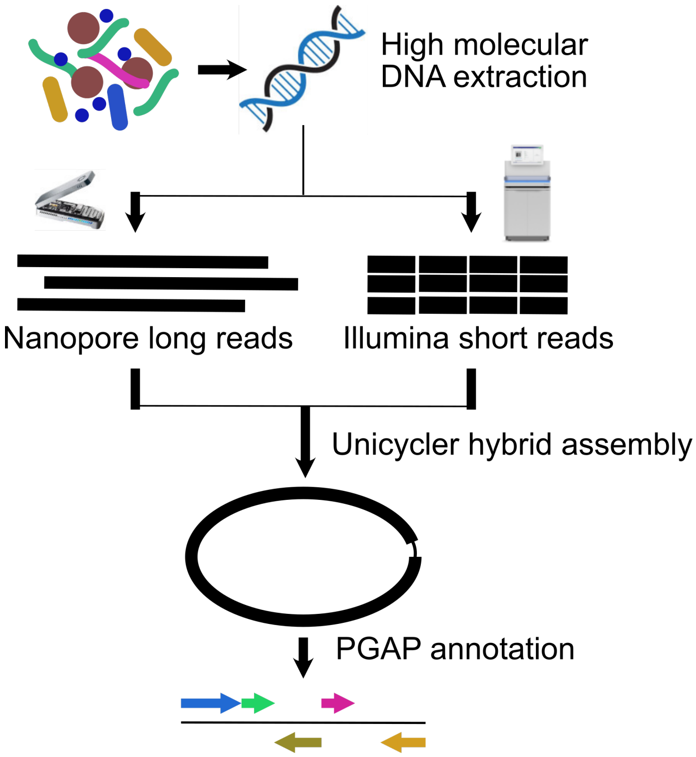


Figure S1: Schematic of hybrid assembly approach.

### Synthetic community strains

Table S1 lists information for all 123 strains in our community. This community is closely modeled after hCom1 and hCom2 from Cheng *et al.* 2021 [4]. We include all strains from both hCom1 and hCom2 in order to start with a comprehensive community. We also include several additional strains (*Peptostreptococcus anaerobius* ATCC 27337, *Peptostreptococcus sp.* CC14N HM 1051, *Clostridium sp.* D5, *Turicibacter sanguinis* DSM 14220) that are known to produce important metabolites *in vivo* [5–7]. In addition, we exclude *Bacteroides rodentium* DSM 26882 because it is not a strain isolated from the human gut. Media and inoculation order describe growth of strains as isolates prior to community assembly. Slow growing fastidious strains are inoculated first 3 days prior to community assembly, fastest growing strains are inoculated last 1 day prior to assembly, and intermediate strains are inoculated 2 days prior. Taxonomic classification obtained by running de novo genomes through GTDB-tk [8]. We manually curate closest available NCBI genomes using strain identifier keyword search, prioritizing complete genomes when multiple entries are present. Closest ANI matches to representative species in UHGG database [9] determined using FastANI [10], which was also used to determine ANI with closest available NCBI genome. Kofamscan [11] was used to obtain KO mappings for both de novo and closest NCBI genome, differential KOs (diffKOs) are counted as those for which the maximum hit bitscore in the de novo genome is more than 2-fold different from the NCBI genome.

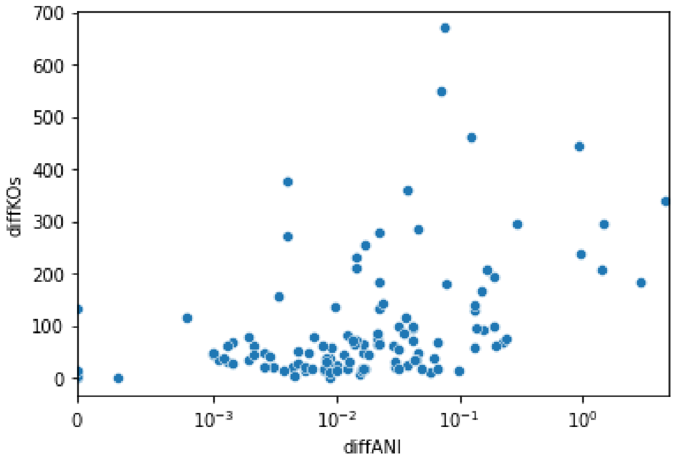


Figure S2: diffANI (100%-ANI) plotted against diffKO plotted for all 123 community strains, comparing de novo vs. NCBI closest genomes.

### Anaerobic chamber and liquid handling setup

To enable large scale liquid handling required for fast and error-free community assembly from isolate strain cultures, we build a custom anaerobic chamber infrastructure that includes liquid handling capabilities in close proximity to culture incubator space and plate reader. Brian please fill this out with any more details you'd like to mention.

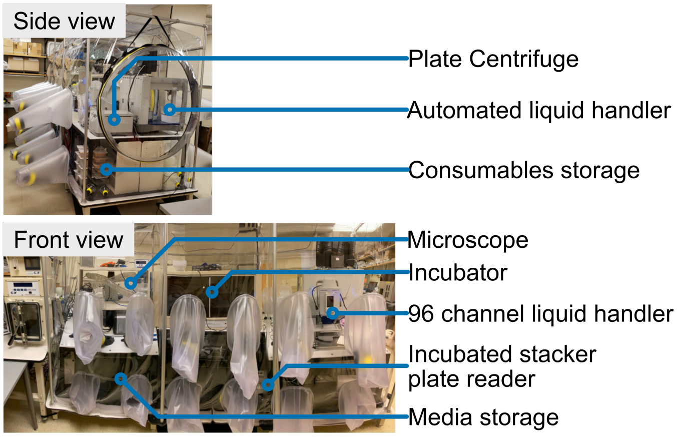


Figure S3: Labeled picture of anaerobic chamber system used for experiments.

### DNA extraction and metagenomic sequencing libraries

A total of 276 DNA samples are extracted (2 sampling × 3 sequencing replicates for inoculum, plus 6 passages × 5 experimental conditions (mucin agar microcosms, mucin agar supernatant, plain agar microcosms, plain agar supernatant, no-microcosm control) × 3 biological replicates (separate culture tubes, i.e., R1, R2, R3) × 3 technical replicates (within culture tubes, e.g., R1a, R1b, R1c). Across samples, we measure mean DNA concentration $7.3ng/uL$ – read libraries generated from these samples after library prep and shotgun metagenomic sequencing exhibit mean read depth $1.2 \times 10^7$. Table S2 details abundance (read fraction) and horizontal coverage for each strain for each read library. Table S3 details log microcosm enrichment scores from paired microcosm-vs.-supernatant samples, as well as aggregated microcosm enrichment scores for both plain-agar and mucin-agar microcosm cultures.

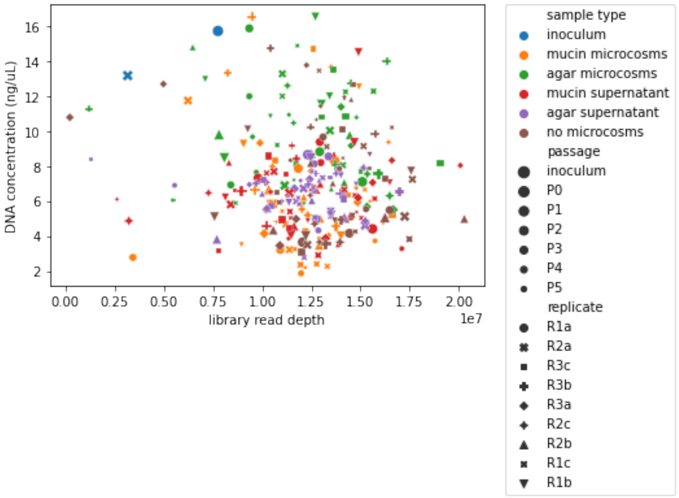


Figure S4: DNA concentration plotted against library read depth for each sample.

### Comparing abundance quantification using Ninjamap with custom database against Kraken2 with UHGG database

To validate the read abundances determined using Ninjamap with our custom strain database, we use the same read libraries and quantify read abundance using Kraken2 [12] with an existing gut species database – UHGG [9]. For each strain, we compare the relative abundance calculated by NinjaMap with that of the closest UHGG species determined by Kraken2. Note that for several strains, there is not a 1-1 correspondence between strain and UHGG species as multiple strains all have the same closest UHGG species (maximum strain correspondence up to 4). In these cases, we sum the NinjaMap relative abundances of all corresponding strains prior to comparing against Kraken2-UHGG species abundance. Across all passage / experimental condition / replicate samples (270 total), we find strong correlation between NinjaMap and Kraken2 relative abundances (min,mean,max $R^2$ of 0.944,0.978 and 0.995 respectively).

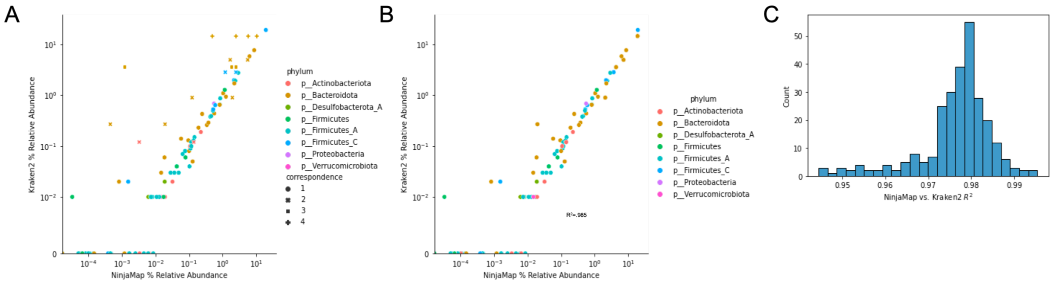


Figure S5: Comparison between relative abundances determined by NinjaMap with custom de novo genomes database versus Kraken2 with UHGG database. **A:** Scatterplot of NinjaMap vs. Kraken2 relative abundances for each strain, mapped to closest UHGG species, using the no-microcosm control passage 1 replicate 1a sample library. Most strains have a 1-1 correspondence to closest UHGG species, but certain species have up to 4 corresponding strains. **B:** Scatterplot of NinjaMap vs. Kraken2 relative abundances for each strain, mapped to closest UHGG species, summing NinjaMap abundances in cases where species have up multiple corresponding strains. **C:** Distribution of $R^2$ values for NinjaMap vs. Kraken2 relative abundances across all passage / experimental condition / replicate samples (270 total).

### Community diversity subset by phylum, and also comparing plain-agar microcosms

Counting detected strains (1% horizontal coverage and 0.01% relative abundance cutoff) subset by phylum, we demonstrate that most of diversity gain in our synthetic community due to microcosm addition occurs in Bacteroides, Firmicutes_A, and Firmicutes. We also plot counts for plain-agar microcosm / supernatant samples, showing that this abundance increase occurs when plain-agar microcosms are added, though to a lesser extent than with mucin-agar microcosms. Finally, we include results from ANOVA with post-hoc Tukey HSD test for significance, comparing strain counts from late passage (P3-P6) no-microcosm, mucin microcosm, mucin supernatant samples. In addition to these 3 experimental conditions we also include a 4th pseudo-condition – mucin readsmix – where NinjaMap analysis is done using pooled mucin microcosm and mucin supernatant reads (each downsampled at 50% to adjust for total read number) from the same culture tube. Regardless of how mucin microcosm cultures are sampled (on microcosms, in supernatant, or pseudomix of reads from both), these cultures exhibit significantly more ($p < 1^{-5}$) detected strains (i.e., community richness) than no-microcosm cultures.

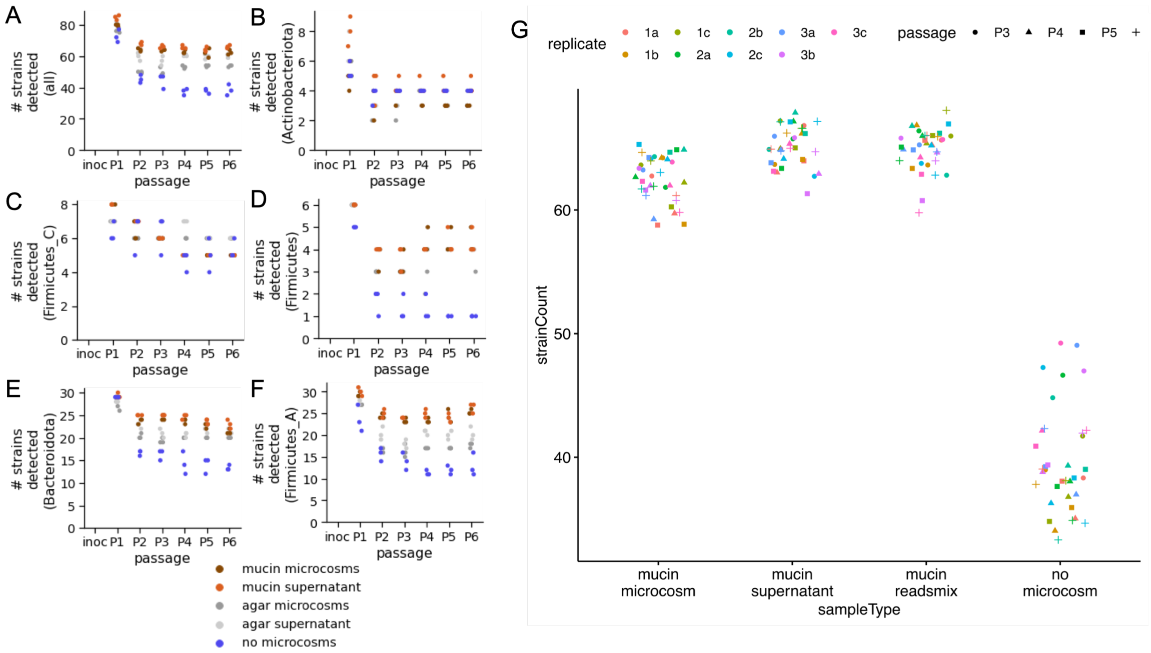


Figure S6: Detected (1% horizontal coverage and 0.01% relative abundance cutoff) strain counts across passages, subset by phylum, including data from mucin-agar microcosm/supernatant, plain-agar microcosm/supernatant, and no-microcosm samples. **A:** Considering all phyla, strain counts are higher when microcosms are present, slightly lower with plain-agar microcosms than mucin-agar. **B:** Actinobacteria does not exhibit strong diversity gain with addition of microcosms. **C:** Firmicutes_C (∼Negativicutes) does not exhibit strong diversity gain with addition of microcosms. **D:** Firmicutes (∼Bacillus) exhibits strong diversity gain with addition of microcosms. **E:** Bacteroidota exhibits strong diversity gain with addition of microcosms. **F:** Firmicutes_A (∼Clostridia) exhibits strong diversity gain with addition of microcosms, particularly mucin-agar. **G:** Comparing all late-passage datapoints (passages P3-P6, biological replicates 1-3, technical replicates a-c, total n=36) for mucin microcosm, mucin supernatant, mucin readsmix and no microcosm conditions. Strain counts in no-microcosm cultures are significantly lower than all mucin conditions. Considering only final passage P6, and collapsing technical replicates into their median value for each biological replicate (i.e., n=3), ANOVA comparison between four conditions yields overall $p = 5.3 \times 10^{-7}$, with pairwise post-hoc Tukey HSD $p = 2.74 \times 10^{-6}, 9.09 \times 10^{-7}, 1.35 \times 10^{-6}$ between no-microcosms versus mucin microcosms, mucin supernatant, and mucin readsmix respectively. Pairwise post-hoc Tukey HSD $p > 0.05$ between all mucin conditions.

### Supplemental abundance measurements with plain agar, highlighting additional strains

We plot here additional heat maps and graphs that include abundance data from the experiment, including data for plain agar microcosm cultures (microcosm and supernatant), for the full 123 community (including less prevalent strains). We also provide additional examples of abundance patterns between related strains.

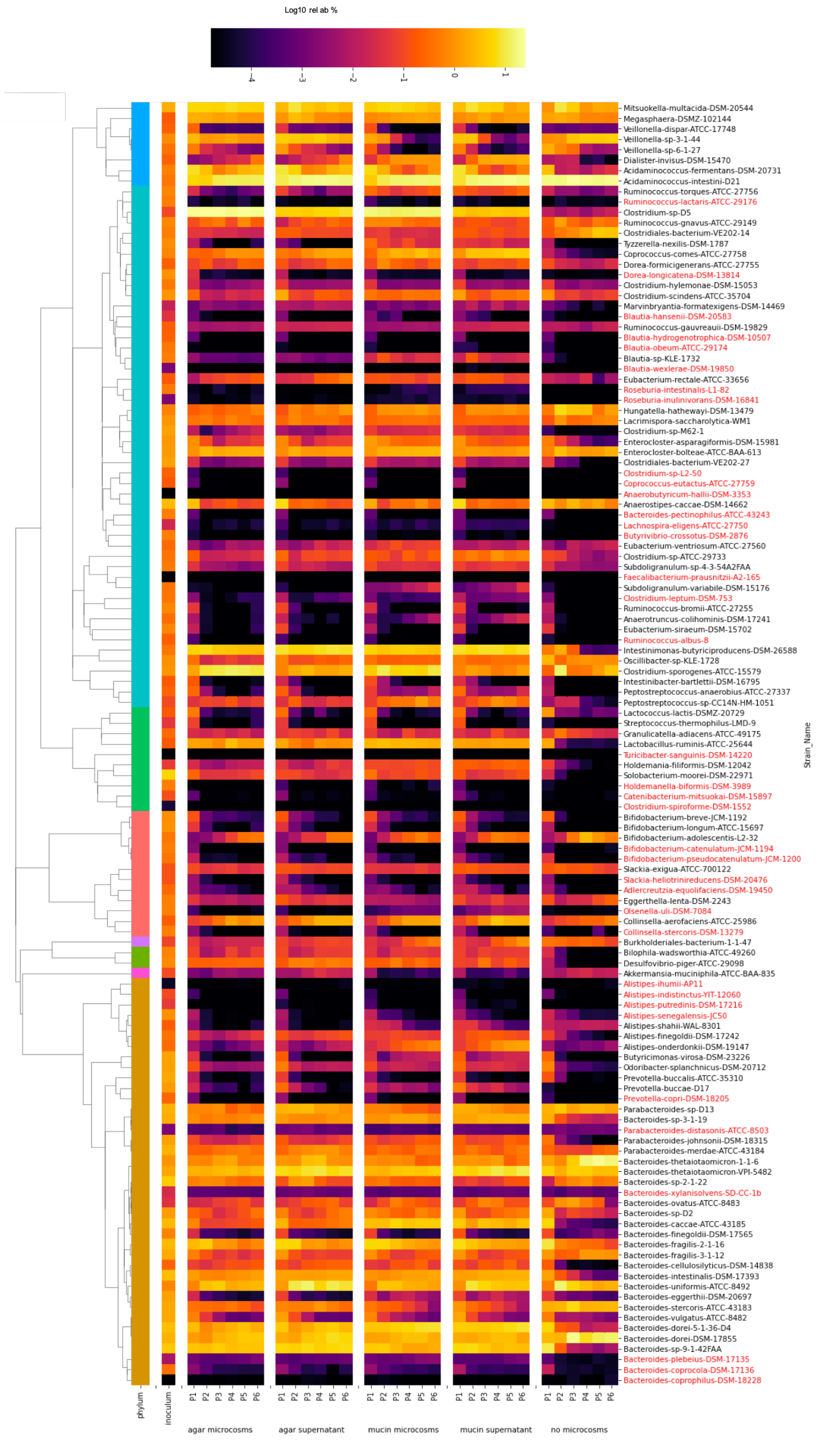


Figure S7: All abundances from across all 5 experimental conditions (mucin agar microcosm/supernatant, plain agar microcosm/supernatant, no microcosm control) and 6 passages plotted as heatmap, taking median of 3 biological replicates, which are themselves median of 3 technical replicates. Strains highlighted in red were not in the top 86 most prevalent strains and were not included in the analysis described in the main text. Strains with low abundance in inoculum are those which failed to grow from glycerol stock isolates.)

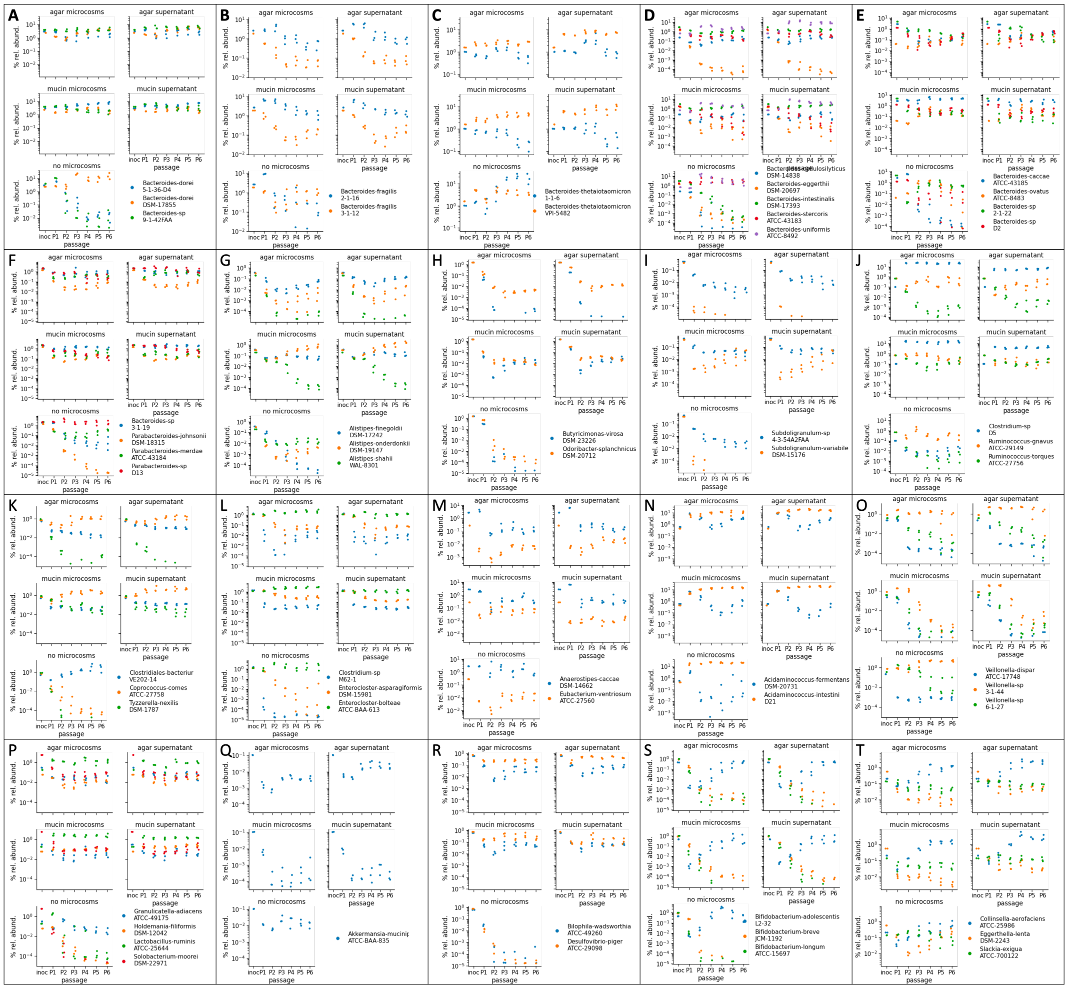
A
B
C
D
E
F
G
H
I
J
K
L
M
N
O
P
Q
R
S
T
agar microcosms
agar supernatant
mucin microcosms
mucin supernatant
no microcosms
% rel. abund.
passage
inoc P1 P2 P3 P4 P5 P6
Bacteroides-dorei 5-1-36-D4
Bacteroides-dorei DSM-17855
Bacteroides-sp 9-1-42FAA
Bacteroides-fragilis 2-1-16
Bacteroides-fragilis 3-1-12
Bacteroides-thetaiotaomicron 1-1-6
Bacteroides-thetaiotaomicron VPI-5482
Bacteroides-cellulosilyticus DSM-14838
Bacteroides-eggerthii DSM-20697
Bacteroides-intestinalis DSM-17393
Bacteroides-stercoris ATCC-43183
Bacteroides-uniformis ATCC-8492
Bacteroides-caccae ATCC-43185
Bacteroides-ovatus ATCC-8483
Bacteroides-sp 2-1-22
Bacteroides-sp D2
Bacteroides-sp 3-1-19
Parabacteroides-johnsonii DSM-18315
Parabacteroides-merdae ATCC-43184
Parabacteroides-sp D13
Alistipes-finegoldii DSM-17242
Alistipes-onderdonkii DSM-19147
Alistipes-shahii WAL-8301
Butyricimonas-virosa DSM-23226
Odoribacter-splanchnicus DSM-20712
Subdoligranulum-sp 4-3-54A2FAA
Subdoligranulum-variabile DSM-15176
Clostridium-sp D5
Ruminococcus-gnavus ATCC-29149
Ruminococcus-torques ATCC-27756
Clostridiales-bacterium VE202-14
Coprococcus-comes ATCC-27758
Tyzzerella-nexilis DSM-1787
Clostridium-sp M62-1
Enterocloster-asparagiformis DSM-15981
Enterocloster-bolteae ATCC-BAA-613
Anaerostipes-caccae DSM-14662
Eubacterium-ventriosum ATCC-27560
Acidaminococcus-fermentans DSM-20731
Acidaminococcus-intestini D21
Veillonella-dispar ATCC-17748
Veillonella-sp 3-1-44
Veillonella-sp 6-1-27
Granulicatella-adiacens ATCC-49175
Holdemania-filiformis DSM-12042
Lactobacillus-ruminis ATCC-25644
Solobacterium-moorei DSM-22971
Akkermansia-muciniq ATCC-BAA-835
Bilophila-wadsworthia ATCC-49260
Desulfovibrio-piger ATCC-29098
Bifidobacterium-adolescentis L2-32
Bifidobacterium-breve JCM-1192
Bifidobacterium-longum ATCC-15697
Collinsella-aerofaciens ATCC-25986
Eggerthella-lenta DSM-2243
Slackia-exigua ATCC-700122

Figure S8: **(Previous page)** Selected abundance comparisons between related strains, across all 5 experimental conditions (mucin agar microcosm/supernatant, plain agar microcosm/supernatant, no microcosm control) and 6 passages. 3 biological replicates points plotted, each point represents median of 3 technical replicates. **A:** 3 B. dorei strains show coexistence with microcosms, and coexistence occurs with both plain-agar and mucin-agar microcosms (extension of Fig. 1F) **B:** 2 B. fragilis strains show increased abundance of one (2-1-16) relative to the other (3-1-12) upon addition of microcosms. **C:** 2 B. thetaiotaomicron strains show one strain (1-1-6) with reduced abundance while the other (VPI-5482) is unaffected upon addition of microcosms. **D:** 5 Bacteroides strains show greater coexistence with microcosms, with B. cellulosilyticus and B. intestinalis in particular exhibiting higher abundance with microcosms. B. eggerthii exhibits this to a lesser extent, only with mucin-agar microcosms, which appears to coincide with decreased abundance of stercoris. B. uniformis remains highly abundant in all conditions. **E:** 4 additional Bacteroides strains show coexistence with microcosms, with B. caccae in particular exhibiting higher abundance with microcosms. **F:** 4 Parabacteroides strains show coexistence with microcosms, with P. johnsonii in particular exhibiting higher abundance with microcosms. **G:** 3 Alistipes strains show higher abundance with microcosms of A. finegoldii and A. onderkii (particularly mucin-agar). These increases appear to come at the expense of A. shahii, which is the most abundant Alistipes strain without microcosms, but least abundant with microcosms. **H:** 2 Odoribacter strains show neither grow well without microcosms – growth is rescued for both in the presence of mucin-agar microcosms, but in plain-agar microcosms culture, only O. splanchnicus is rescued. **I:** 2 Subdoligranulum strains show coexistence with mucin-agar microcosms, but not plain-agar microcosms (extension of Fig. 1G) **J:** 3 Lachnospiraceae strains show distinct responses to presence of microcosms: sp. D5 exhibits improved growth in the presence of either mucin agar or plain agar microcosms, R. torques only for mucin agar microcosms, while R. gnavus does not exhibit large abundance changes to microcosm presence. **K:** 3 additional Lachnospiraceae strains shows coexistence with microcosms. Only strain VE202-14 is abundant without microcosms, but C. comes in particular comes more abundant with mucin-agar and plain-agar microcosms added, while T. nexilis growth is rescued with mucin-agar microcosms only. **L:** 3 additional Lachnospiraceae strains shows coexistence with microcosms, while E. bolteae dominates among the three without microcosms **M:** 3 additional Lachnospiraceae strains shows approximately similar abundances with and without microcosms. **N:** 2 Acidaminococcus strains show coexistence with microcosms, and coexistence occurs with both plain-agar and mucin-agar microcosms (extension of Fig. 1H) **O:** 3 Veillonella strains show reduced abundance upon addition of mucin agar microcosms, particularly for sp. 3-1-44 – this effect is less pronounced for plain-agar microcosms. **P:** 4 Bacillus strains shows higher abundances with microcosms, particularly for L. ruminis. **Q:** Akkermansia muciniphila ATCC-BAA-835 show lower abundance when mucin-agar microcosms are present, compared with both plain-agar microcosms and no-microcosm control. **R:** 2 Desulfobacter strains show both have higher growth when microcosms are present. **S:** 3 Bifidobacterium strains show patterns which are not strongly affected by microcosms (B. adolescentis dominates in all conditions). **T:** 3 Coriobacteria strains show distinct responses to presence of microcosms: C. aerofaciens exhibits improved growth in the presence of microcosms, E. lenta exhibits reduced growth, while S. exigua appears relatively unaffected by comparison.

### Microcosm enrichment with plain agar

Here we show microcosm enrichment calculations with the plain agar microcosms, both as heatmap of individual strain as well as aggregated by phylum. We also highlight some additional enrichment patterns between strains.

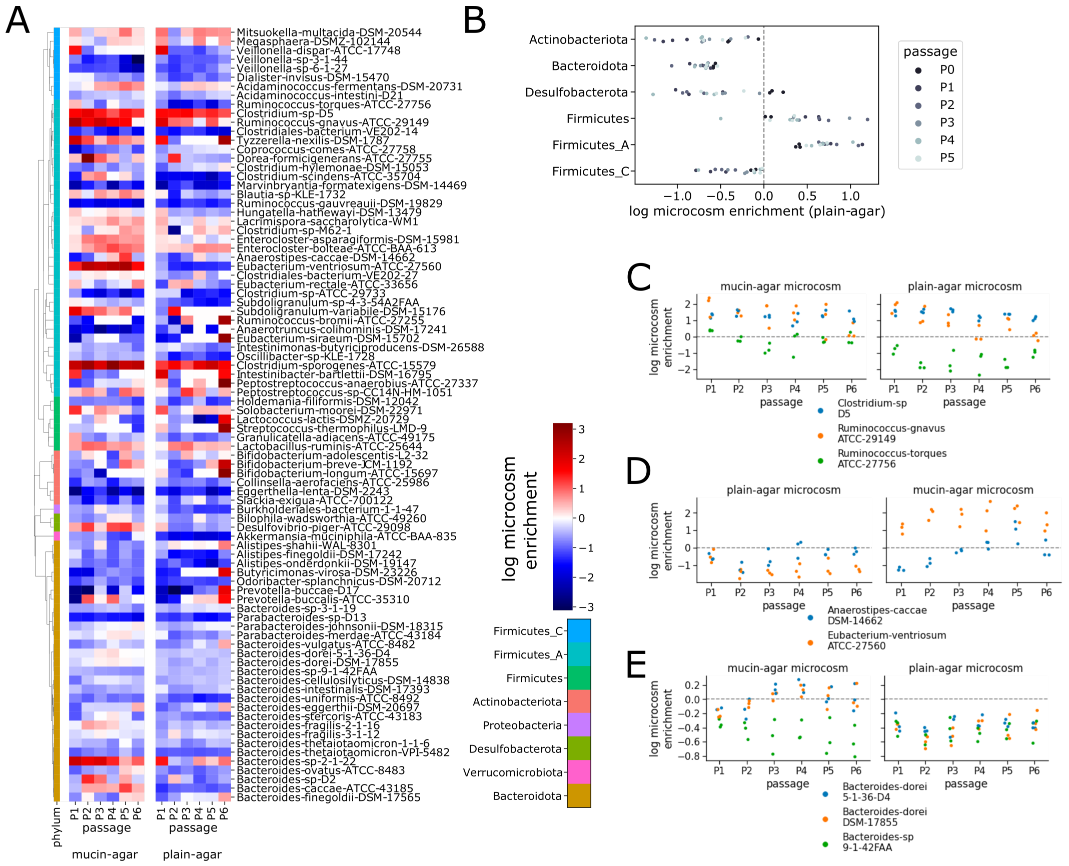


Figure S9: Comparison between microcosm enrichment with mucin-agar and plain-agar microcosms. **A:** Heatmap of mucin-agar (left, copied from Fig. 2A) and plain-agar microcosm enrichment scores – positive (red) scores indicate microcosm enrichment. **B:** Aggregated at phylum level, results are similar in plain-agar microcosm cultures as mucin-agar with the exception of Desulfobacterota, which is now enriched in supernatant (compare with Fig. 2B). **C:** Clostridium sp. D5 exhibits microcosm enrichment for both mucin-agar and plain-agar. **D:** Eubacterium ventriosum exhibits microcosm enrichment for mucin-agar only. **E:** All three strains of B. dorei exhibit microcosm depletion in plain-agar case.

### Gene difference comparisons between strains

We search for differential KO / gene family presence between strain genomes by applying a 1.5-fold hmmer bitscore cutoff for each KO. For example, if a strain has a maximum bitscore of 10 for a particular KO, any strain with a maximum bitscore greater than 15 or less than 6.67 will be considered to have a differential KO presence. Table S4 lists maximum bitscore hits for K00441 and K08217 KO pHMMs against Bacteroides dorei DSM-17855 / Bacteroides dorei 5-1-36-D4 / Bacteroides sp. 9-1-42FAA, as well as the maximum bitscore hits for K14440 in Subdoligranulum variabile-DSM-15176 / Subdoligranulum-sp-4-3-54A2FAA, and K14743 in Acidaminococcus fermentans DSM-20731 / Acidaminococcus intestini D21, supporting the examples from Fig. 2C-E.

### Gene neighbourhood analysis of K00441

Table S5 lists frequency of co-occurrence between every KO and K00441 (number of instances they exist within 10kb of each other, across all community genomes), as well as the number of times out of 1000 random permutations (all gene labels shuffled across all genomes) that the actual frequency exceeds the random permutation.

### *In vivo* dataset analysis

We use the Suez *et al.* 2018 dataset [13] as a source of metagenomic read libraries from *in vivo* gut microbial samples, with paired lumen and mucosa samples within individuals, at multiple gastrointestinal tract locations. We initially select 16 individuals with lumen and mucosa data from terminal ileum, cecum and ascending colon. We then use Kneaddata (part of Biobakery suite []) to filter reads based on quality and to remove human (host) reads. After this filtering step, 13/16 individuals remain, corresponding to a total of 78 read libraries – Table S6 details read libraries used.

We then use Kraken2 [12] to classify filtered reads from each of these libraries against the UHGG database [9], obtaining phyla-level and species-level relative abundance estimates for each read library. We focus our analysis on species that are detected with at least 0.01% abundance in at least 10% of libraries, leading to a subset of 676 species. For each of these species, we calculate their mucosal enrichment score by comparing paired mucosa/lumen samples from the same individual and site, applying the log-ratio approach as done for microcosm enrichment with the in-vitro dataset. We additionally generate an aggregate score by taking the mean over standard deviation of log-ratios. Aggregate scores are generated both per individual (over 3 sites), as well as across all individuals to generate a single aggregate score per species. These enrichment scores are detailed in Table S7. We also calculated phyla-level enrichment scores for each paired lumen/mucosa sample, plotted in Fig. S10A. Similar trends exist between *in vivo* and *in vitro* enrichments at phylum level: Bacteroidota is enriched toward both supernatant (*in vitro*) and lumen (*in vivo*), while Firmicutes_A (Clostridia-like) and Firmicutes (Bacillus-like) are enriched toward microcosm / mucosa. However, discrepancies also exist, as Actinobacteriota is enriched toward supernatant *in vitro* and mucosa *in vivo*.

For the species-level results, we compare the spearman correlation of *in vivo* mucosal enrichment scores across species with *in vitro* microcosm enrichment scores (both mucin-agar and plain agar) across strains, mapping *in vitro* strains to their closest UHGG species and only considering taxa that pass the prevalence threshold (0.01%

abundance in at least 10% of samples) in both analyses. We find significant positive pairwise Spearman correlation scores between *in vivo* mucosal-enrichment and *in vitro* microcosm-enrichment scores using plain agar microcosms ($p<0.001$), as well as *in vitro* plain-agar microcosms and mucin microcosms scores ($p<0.001$). Correlation is positive but not significant ($p=0.16$) between *in vivo* and *in vitro* mucin microcosms. Stratifying the *in vivo* dataset by human participants, we observe significant variability between participants, who group into two main clusters. Our *in vitro* log-microcosm-enrichment scores (plain-agar and mucin-agar microcosms) group within the larger of these two clusters, indicating that observed discrepancy between *in vitro* and *in vivo* mucosal/microcosm scores does not exceed inter-subject variability.

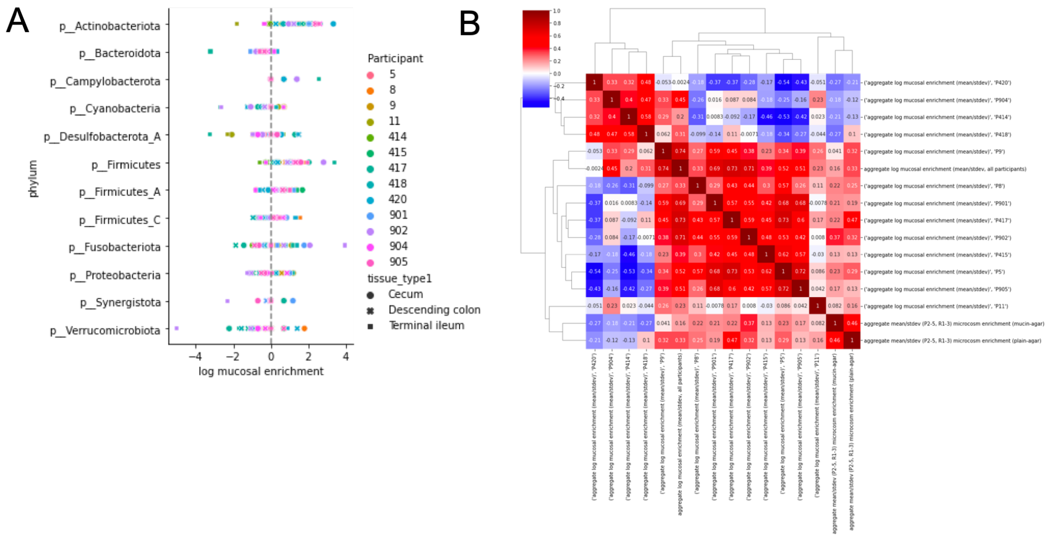


Figure S10: Mucosal enrichments from *in vivo* data [13]. **A:** Mucosal enrichments from *in vivo* dataset at phylum level. **B:** Spearman correlation of mucosal enrichments at species-level, compared with *in vitro* strain microcosm enrichmments.

### Detailed results from phylogenetic linear models

Table S8 lists all phylogenetic linear model results for all tested KOs. For the 244 microcosm-associated KOs (i.e., effect size $> 0$ ) that pass Benjamini-Hochberg FDR significance test at $p=0.01$. We also show clade specific results in Table S9, where only the subset of strains belonging to a single clade are considered when performing the the phylogenetic linear model – note FDR requirement is relaxed for clade specific significance testing to increase sensitivity. We group phylum Firmicutes, Firmicutes_A and Firmicutes_C into a single clade for this analysis. The majority of significant phyla-specific hits (217) occur within Bacteroidota – which has numerous instances of closely related strains – followed by Firmicutes (5); other phyla with fewer strain representatives did not produce clade-specific significant hits. We also include results from parallel analysis using the Suez 2018 *in vivo* dataset (Table S10), where we identify a total of 6831 significant KO hits associated with increased mucosal enrichment. Comparing the *in vitro* and *in vivo* KO hits (Fig. S11), we find that 199 KOs are associated with both increased mucosal (*in vivo* dataset [13]) and microcosm (*in vitro* dataset from our own work) enrichment, 45 that are only microcosm-enrichment associated, 6632 that are only mucosal-enrichment associated, and 5984 that are neither, corresponding to a log odds ratio of 3.99 and $p < 1^{-20}$ using Fisher's exact test, confirming significant overlap between *in vivo* and *in vitro* results.

| In vivo data (Suez 2018) | In vitro data (this work): Significant microcosm enriched | Not significant microcosm enriched |
|---|---|---|
| Significant mucosa enriched | 199 | 45 |
| Not significant mucosa enriched | 6632 | 5984 |

Figure S11: 2x2 contingency table showing overlap between significant KO hits for microcosm enrichment (*In vitro* data from this work) and significant KO hits for mucosa enrichment (*In vivo* data from Suez 2018 [13]). Log odds ratio of 3.99 and $p < 1^{-20}$ using a 2x2 fisher exact test.

### Aggregative KEGG BRITE analysis

For each KEGG BRITE hierarchical category, we consider all KOs that full under the category's umbrella. We then intersect this subset of BRITE KO's with the 244 significantly microcosm-associated KOs identified earlier, to generate a 2x2 contingency table (microcosm-associated KOs in BRITE category, non-microcosm-associated KOs in BRITE category, microcosm-associated KOs not in BRITE category, and non-microcosm-associated KOs not in BRITE category). We then use this table to calculate log odds ratio p values using a fisher exact test – this yields a total of 43 BRITE categories significantly ($p < 0.05$) associated with increased microcosm enrichment, listed in Table S11.

### Identifying and grouping biosynthetic gene clusters using DeepBGC and hierarchical clustering

We use DeepBGC [14] to identify BGCs in our genomes, yielding a total of 1349 BGCs in our community genomes. For each BGC, we determine KEGG KO presence by considering all CDSs within the BGC – all KO's mapped to any CDS within the BGC (at least 0.5x hmmer bitscore relative to the KEGG-defined bitscore threshold, and 0.5x length overlap relative to total pHMM length) are considered present in the BGC. Based on this we focus on 1103 BGCs with at least 3 mapped KOs (and conversely KOs with at least 3 mapped BGCs), yielding a boolean 1103 BGC x 1387 KO presence absence matrix. We apply hierarchical clustering on this matrix using Jaccard distance metric to generate 256 groups of BGCs (see Table S12), and subsequently map each BGC back to its original strain. This finally yields a boolean 86 strain x 256 BGC-group presence absence matrix (again focusing on the 86 most prevalent strains, see Fig. S12). We then test iteratively using phylolm [15] one BGC-group at a time for significant association with microcosm enrichment across strains. Note that there are instances where numerous BGC-groups exhibit very similar genotypes across these 86 strains, leading to the case where a number of BGC-groups exhibit similar p-values (see Fig. 5B). Table S13 lists the all KOs in the 7 significantly microcosm-enriched BGC-groups.

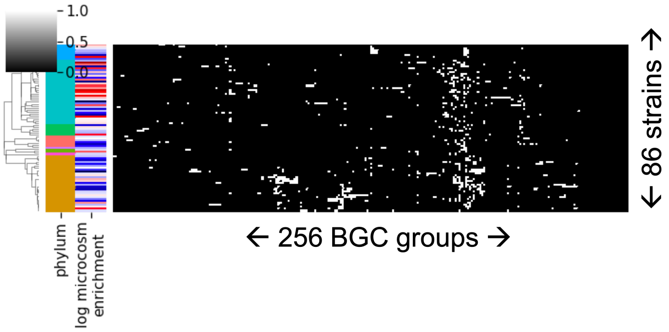


Figure S12: BGC-group by strain presence absence matrix. White indicates BGC-group is present in strain, phylum colors as in Fig. 1.

[6] Julia H Kemis, Vanessa Linke, Kelsey L Barrett, Frederick J Boehm, Lindsay L Traeger, Mark P Keller, Mary E Rabaglia, Kathryn L Schueler, Donald S Stapleton, Daniel M Gatti, Gary A Churchill, Daniel Amador-Noguez, Jason D Russell, Brian S Yandell, Karl W Broman, Joshua J Coon, Alan D Attie, and Federico E Rey. Genetic

determinants of gut microbiota composition and bile acid profiles in mice. *PLOS Genetics*, 15(8):e1008073, aug 2019.
